# *RFC1* regulates the expansion of neural progenitors in the developing zebrafish cerebellum

**DOI:** 10.1101/2024.06.28.601164

**Authors:** Fanny Nobilleau, Sébastien Audet, Sanaa Turk, Charlotte Zaouter, Meijiang Liao, Nicolas Pilon, Martine Tétreault, Shunmoogum A. Patten, Éric Samarut

**Affiliations:** Department of Neuroscience, Faculty of Medicine, Université de Montréal, Montreal, H3T 1J4, Quebec Canada; Functional Genomics of Rare Diseases Laboratory, Research Center of the University of Montréal Hospital Center (CRCHUM), Montreal H2X 0A9, Quebec Canada; INRS-Centre Armand-Frappier Santé Biotechnologie, Laval, H7V 1B7, Quebec Canada; Molecular Genetics of Development Laboratory, Département des Sciences Biologiques, Université du Québec à Montréal (UQAM), Montréal H3C 3P8, Quebec Canada; Centre d’excellence en recherche sur les maladies orphelines – Fondation Courtois (CERMO-FC), Université du Québec à Montréal, Montreal H2X 3Y7, Quebec Canada; Department of Pediatrics, Faculty of Medicine, Université de Montréal, Montreal, 1C5, Quebec Canada

**Keywords:** Keywords, RFC1, cerebellum, neurogenesis, zebrafish, late-onset cerebellar ataxia, CANVAS

## Abstract

DNA replication and repair are basic yet essential molecular processes for all cells. *RFC1* encodes the largest subunit of the Replication Factor C (RFC), which is a clamp-loader during DNA replication and repair. Intronic repeat expansion in *RFC1* has recently been associated with so-called *RFC1*-related disorders, which mainly encompass late-onset cerebellar ataxias. However, the mechanisms that make certain tissues more susceptible to defects in these universal pathways remain mysterious. In this study, we provide the first investigation of *RFC1* gene function *in vivo* using zebrafish. We showed that *RFC1* is expressed in neural progenitor cells within the developing cerebellum and that it is necessary to maintain these cells’ genomic integrity during neurogenic maturation. Accordingly, *RFC1* loss-of-function leads to a severe cerebellar phenotype due to impaired neurogenesis of both Purkinje and granule cells. Our data thus point to a specific role of *RFC1* in the developing cerebellum, paving the way for a better understanding of the pathogenic mechanisms underlying *RFC1*-related disorders.

## Introduction

The cerebellum is the main derivative of the embryonic hindbrain in all vertebrates. It receives and processes information from sensory systems, the spinal cord and other parts of the brain ^1-4^, using these inputs to ensure that motor movements are smooth, coordinated and efficient^5^. Best known for this motor coordination role, the cerebellum also influences cognitive processes such as attention, language, and modulation of emotional responses ^6-8^. Due to all these essential roles, dysfunctions in the development or activity of the cerebellum lead to pathological syndromes associated with movement disorders, problems of balance, posture, and even learning ^8-10^.

Ataxias are neurodegenerative diseases of the cerebellum and/or brainstem that are characterized by a lack of muscle control or coordination of voluntary movements. These conditions have been linked to hundreds of genes with high phenotypic overlap ^11,12^. Particularly, late-onset cerebellar ataxias (LOCA) are a heterogeneous group of cerebellar progressive neurodegenerative diseases that manifest in adulthood with unsteadiness. It is estimated that up to 3 in 100,000 people worldwide will develop a LOCA. This prevalence is likely underestimated because of the great heterogeneity of clinical presentations, especially in aging patients. The genetic and phenotypic heterogeneity of LOCA contributes to our inability to diagnose ataxia patients, and most LOCA patients remain without a genetic diagnosis ^13^. This dampens our understanding of the underlying pathological mechanisms involved in these diseases. However, in 2019, a breakthrough in the genetics of ataxia identified biallelic repeat expansions in the second intron of the *RFC1* gene, encoding Replication Factor C subunit 1. These pentanucleotide repeat expansions are primarily associated with the cerebellar ataxia-neuropathy-vestibular areflexia syndrome (CANVAS) ^14,15^. Although it is a rare syndrome, recent findings suggest that *RFC1* expansions are a major cause of a broader phenotypic spectrum of late-onset progressive ataxias. Moreover, the pathogenic scope of these expansions has been expanded to other movement syndromes, now collectively referred to as *RFC1-related disorders*, which include specific forms of parkinsonism and multiple systems atrophy (MSA) ^16-19^. Although the pathogenicity of *RFC1* repeat expansions is now well established, the underlying mechanisms remain unknown. Recent studies propose that *RFC1* loss-of-function could be at the basis of the repeat expansion pathogenicity ^20-23^, as the recessive inheritance pattern also suggests. However, other studies showed no specific neurodegenerative phenotype in *RFC1*-depleted neuronal cultures *in vitro* ^24^. This questions the direct involvement of the *RFC1* gene in so-called *RFC1*-related pathologies.

*RFC1* encodes the largest subunit of the replication factor C, a DNA-dependent ATPase which loads the DNA-clamp protein PCNA (Proliferating Cell Nuclear Antigen) and recruits DNA polymerases onto DNA undergoing replication or repair ^25-27^. The Human Protein Atlas database indicates a ubiquitous tissue expression in adults, which is consistent with its assumed housekeeping function as a DNA repair and replication regulator. Nevertheless, the accurate expression of *RFC1* has not yet been assessed in mammals or vertebrates *in vivo*, particularly during embryogenesis. Since its cloning in humans in the 90s ^28^, the function of *RFC1*, particularly in the context of DNA replication and repair, has only been studied *in vitro* ^25,29,30^ or in plants ^31,32,33^). Detailed *in vivo* analysis of its function in animals is needed to better understand *RFC1*-associated diseases.

In this work, we studied the gene and protein expression patterns of RFC1 in the developing cerebellum of both mice and zebrafish. This analysis revealed specific expression patterns in neuronal progenitor populations of both Purkinje and granule cell lineages. To circumvent the early embryonic lethality of *Rfc1* knockout mouse models, we generated a zebrafish model of *rfc1* loss-of-function using CRISPR/Cas9 targeted mutagenesis. Although *rfc1^-/-^*zebrafish larvae die prematurely after 10 days of age, we found that this does not prevent the functional analysis of RFC1 during neurodevelopment. This model uncovered a key role for *rfc1* in the proper development of both granule and Purkinje cell populations. Notably, single-cell transcriptomic approaches showed that *rfc1* regulates the expansion and differentiation of the neuronal progenitor pools. In the absence of rfc1, these progenitors accumulate DNA damages and stop proliferating, leading to their death and ultimately severely affecting cerebellum neurogenesis. This specific role of RFC1 during neurodevelopment opens new doors regarding its potential involvement in cerebellar pathologies.

## Results

### RFC1 is expressed in the developing cerebellum

To evaluate the tissue distribution of *RFC1 in vivo*, we first evaluated its mRNA expression in zebrafish embryos via whole-mount *in situ* hybridization (Figure 1A). We found an early expression in the developing brain, from 24 hours post-fertilization (hpf), with a specific expression in the developing hindbrain (arrowhead, Figure 1A). At later stages, *rfc1* expression becomes limited to the upper rhombic lip (URL) of the anterior hindbrain, constituting the presumptive cerebellum ^34^. At 120 hours post-fertilization, *rfc1* expression is restricted to the lobus caudalis cerebelli (LCa). Although specific, the expression of *rfc1* in the developing cerebellum is not exclusive, as a signal is also observed in the eye, floor plate and surrounding pharyngeal tissues. To complement these observations, we then examined the RFC1 protein distribution in the developing cerebellum of wild-type mice at the cellular level (Figure 1B). Using co-immunolabelling of RFC1 expression with specific markers of Purkinje cells (PC, Calbindin) or granule cells (GC, Pax6), we showed that RFC1 is expressed in early Purkinje cells progenitors (PCp) from P0. Its expression persists in the Purkinje Cell Layer (PCL) at P7, P11 and P60 (Figure 1B, white arrows). Of note is that the nuclear expression of RFC1 in PCp gradually becomes cytoplasmic in more mature PC from P11 onwards. We also found that RFC1 is not expressed in early granule cell progenitors at P0, as shown by the absence of colocalization with PAX6 marker. However, it is transiently expressed in some migratory granule cells (mGCs) at P11. This stage corresponds to the moment when GCs migrate from the External Granule Layer (EGL) towards the Internal Granular Layer (IGL), passing through the Molecular Layer (ML) and the Purkinje Cell Layer (PCL) (Figure 1B, white arrowheads). Once in the IGL, we could not detect colocalization between RFC1 and the GC marker PAX6. However, we did detect RFC1 expression in some PAX6-positive GCs scattered in the ML. Initially described as ectopic GCs (eGCs), these cells have been recently renamed molecular GCs (mGCs) based on their proven function within the cerebellum circuitry ^35^. Altogether, our results show that RFC1 is expressed in the developing cerebellum, both in PC and transiently in a subset of GCs in vertebrates.

**Figure 1:**
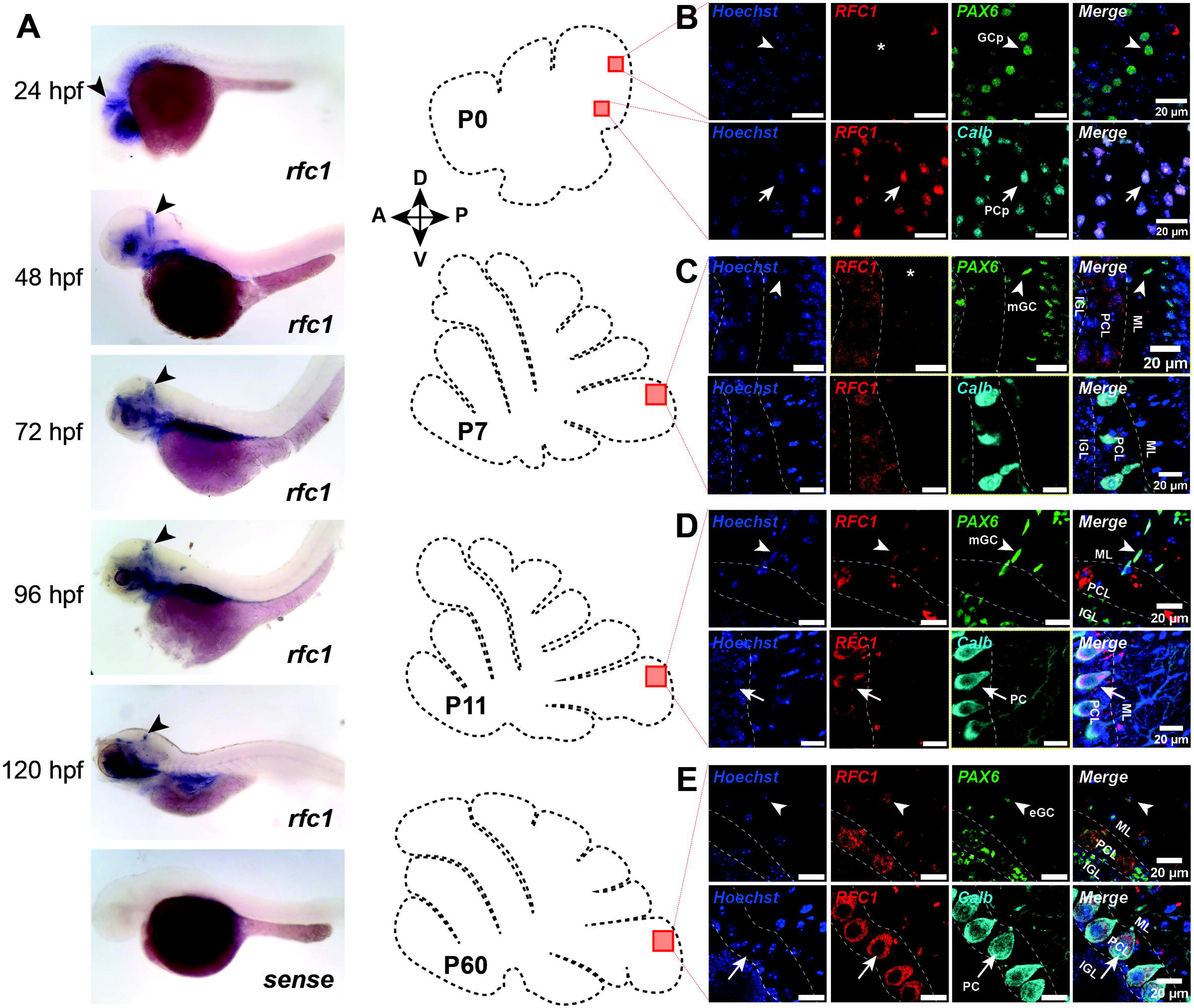
*RFC1* is expressed in the developing cerebellum in the developing cerebellum. (A) Whole-mount in situ hybridization against *rfc1* shows a specific, although not exclusive, expression in the developing cerebellum in zebrafish from 24 hpf (arrowheads). (B, C, D, E) Co-immunofluorescent labelling of RFC1 and PAX6 (granule cell marker) or Calbindin (Calb, Purkinje cell marker) in the developing mouse cerebellum at different stages: P0 (B), P7 (C), P11 (D), P60 (E). RFC1 is expressed in Purkinje Cell progenitors (PCp, arrows in (B)) but not in Granule Cells progenitors (GCp, asterisks and arrowheads in (B)). However, RFC1 is detected in some migratory granule cells from P11 (mGC, arrowheads in (D)) and in Purkinje cells at P7 and P11 (PC, arrows in (C) and (D)). At P60 (E), RFC1 is detectable in some ectopic granule cells from the molecular layer (ML) (eGC, arrowheads in (E)) and in Purkinje cells (PC, arrows in (E)). *ML: Molecular Layer; PCL: Purkinje Cell Layer; IGL: Internal Granular Layer*.

### *Rfc1* loss-of-function leads to premature death, head morphological anomalies and motility defects in zebrafish

The Jackson Laboratory Phenotyping Center generated a knock-out (KO) *Rfc1* allele in mice (*Rfc1*^tm1b(KOMP)Mbp^), for which homozygosity was reported to lead to embryonic and/or pre-weaning lethality (no homozygotes over 66 pups at weaning; MGI ID:5495286 ^36^). In an effort to circumvent this roadblock, we generated a zebrafish *rfc1* KO model using the CRISPR/Cas9 technology. We targeted the 5^th^ exon and selected a founder line with a 20-bp deletion mutation, which leads to a premature stop codon at position 182 in the zebrafish RFC1 protein (Figure 2A, B). Of note is that zebrafish and human RFC1 protein share 57 % of sequence identity (69 % of sequence similarity), encompassing the same functional domains (Figure 2C, Figure S1). We confirmed *rfc1* loss-of-function at the transcript level both in heterozygous (+/-) and homozygous (-/-) zebrafish larvae (Figure 2D). While *rfc1^+/-^* fish survive until adulthood with a normal Mendelian inheritance pattern, we found that *rfc1^-/-^*larvae die prematurely from 8 days of age with no survivor past day 10 (n = 168; Figure 2E). Moreover, *rfc1^-/-^* larvae depict gross morphological abnormalities from 2 days post-fertilization (dpf). In particular, they display a smaller head, reduced eye diameter and a shorter otic vesicle than their siblings (Figure S2A-2B, Figure 2F-2H). These morphological defects become more obvious from 3 dpf and worsen until death. Noteworthy, we did not observe differences in the average body size (Figure S2B), suggesting that the head morphological phenotype observed in *rfc1^-/-^*larvae is very specific and not due to developmental delay. Finally, we monitored the swimming behaviour of 4 dpf larvae and found that *rfc1^-/-^* larvae are significantly hypoactive compared to their siblings as assessed by the total distance swam during 60 min (Figure 2I-2J). Altogether, these data show that RFC1 is necessary for normal development, motor behaviour and survival in zebrafish. Moreover, its loss of function leads to head morphological deformities that we aim to characterize further.

**Figure 2:**
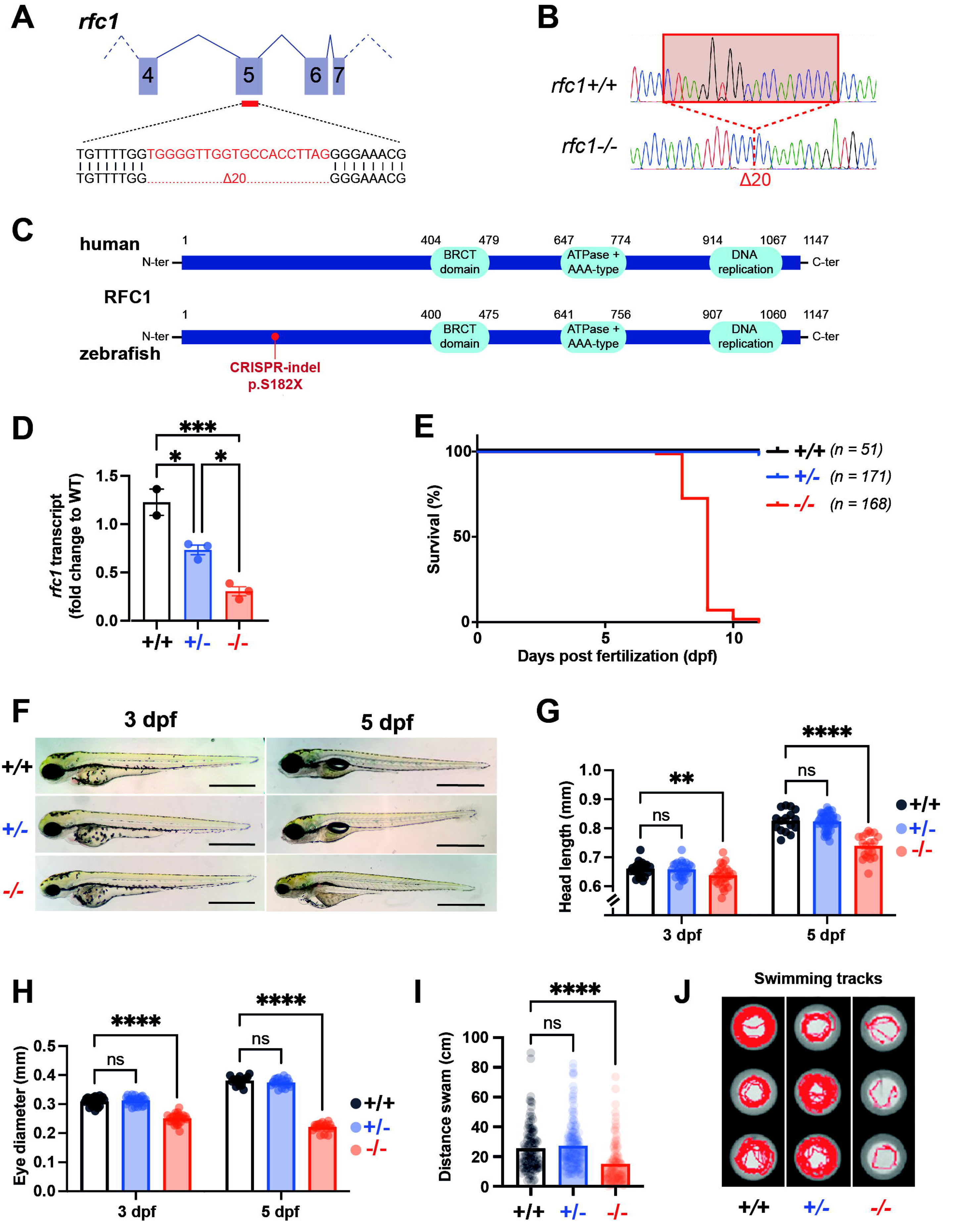
*Rfc1* loss-of-function leads to premature death, head morphological anomalies and motility defects in zebrafish. (A, B) A 20-bp deletion in exon 5 of the zebrafish *rfc1* gene was identified and confirmed by Sanger sequencing (B). (C) This deletion leads to a premature stop codon in position 182 in the zebrafish protein. Of note is that all three RFC1 functional domains (BRCT, ATPase+ AAA-type and DNA replication) are located downstream the premature stop. (D) We confirmed that the level of *rfc1* expression, at the transcript level is significantly reduced in +/- (0.73, ±SEM 0.05) and -/- (0.31, ±SEM 0.05). (5) *rfc1^-/-^* larva dire prematurely by 10 days of age. (F) *rfc1^-/-^* but not *rfc1^+/-^* depict gross morphology defects from 3 dpf characterized by a reduced head and eye size. (G) Quantification of the head length showing a significant reduction in *rfc1^-/-^*larvae from 3 dpf. (H) Quantification of the eye diameter showing a significant reduction in *rfc1^-/-^*larvae from 3 dpf. (I) Quantification of the distance swam for one hour (light off) showing a significant hypomotility of *rfc1^-/-^* larvae compared to siblings (diameter showing a significant reduction in *rfc1^-/-^* larvae from 3 dpf. (J) Swimming tracks over 10 minutes of 5 dpf larvae showing the hypomotility of *rfc1^-/-^* larvae. *ns : P > 0.05; * P* ≤ *0.05; ** P* ≤ *0.01; *** P* ≤ *0.001; **** P* ≤ *0.0001*

### *Rfc1-/-* zebrafish larvae depict severe cerebellar defects

As we showed that *RFC1* is expressed in the developing cerebellum (at the level of the URL in zebrafish and in cerebellar neuronal progenitors and Purkinje Cells in mice, Figure 1), we monitored cerebellum development in our *rfc1^-/-^* zebrafish model. We confirmed that the reduced head size of *rfc1^-/-^* larvae is accompanied by a general reduction in brain size as revealed by hematoxylin-eosin staining of brain cross-sections (Figure 3A). More importantly, unlike in control *rfc1^+/-^*and wildtype larvae, we were not able to identify any distinct cerebellar structures in 4 dpf *rfc1^-/-^*larvae cross-sections at the level of the hindbrain (asterisks in Figure 3A and Figure S3A). To confirm this observation, we followed by *in toto* immunolabelling the development of the two main cerebellar neuronal cell types: Parvalbumin7+ PCs and neurod1+/vglut1+ GCs (Figure 3B-3D, Figure S3B-S3C). Consistent with our histological observations, we showed that GCs fail to develop in *rfc1^-/-^* larvae compared to control siblings (Figure 3B, C). Indeed, the number of neurod1+ GCs is drastically reduced at 3 dpf in *rfc1^-/-^*larvae (Fig. 3B), also remaining low afterwards (Figure 3B, 3E, Figure S3B-S3C). This is consistent with the absence of *vglut1+* posterior axonal projections in *rfc1^-/-^* larvae, which are normally observed from 3 dpf and thicken with age in siblings (Figure 3C, Figure S3C). Similarly, albeit less severe, the number of developing parvalbumin7+ PCs is significantly reduced in *rfc1^-/-^* from 3 dpf onwards (Figure 3D, 3F, Figure S3D). Moreover, the general organization of the PC layer is affected in 5 dpf larvae with the absence of the Valvula cerebelli structure, a cerebellar structure unique to ray-finned fishes that controls non-locomotor related functions ^37^ (Figure 3D, Figure S3D). Overall, these data confirm that *rfc1* is involved in cerebellum development, particularly in populating its neuronal subtypes in vertebrates.

**Figure 3:**
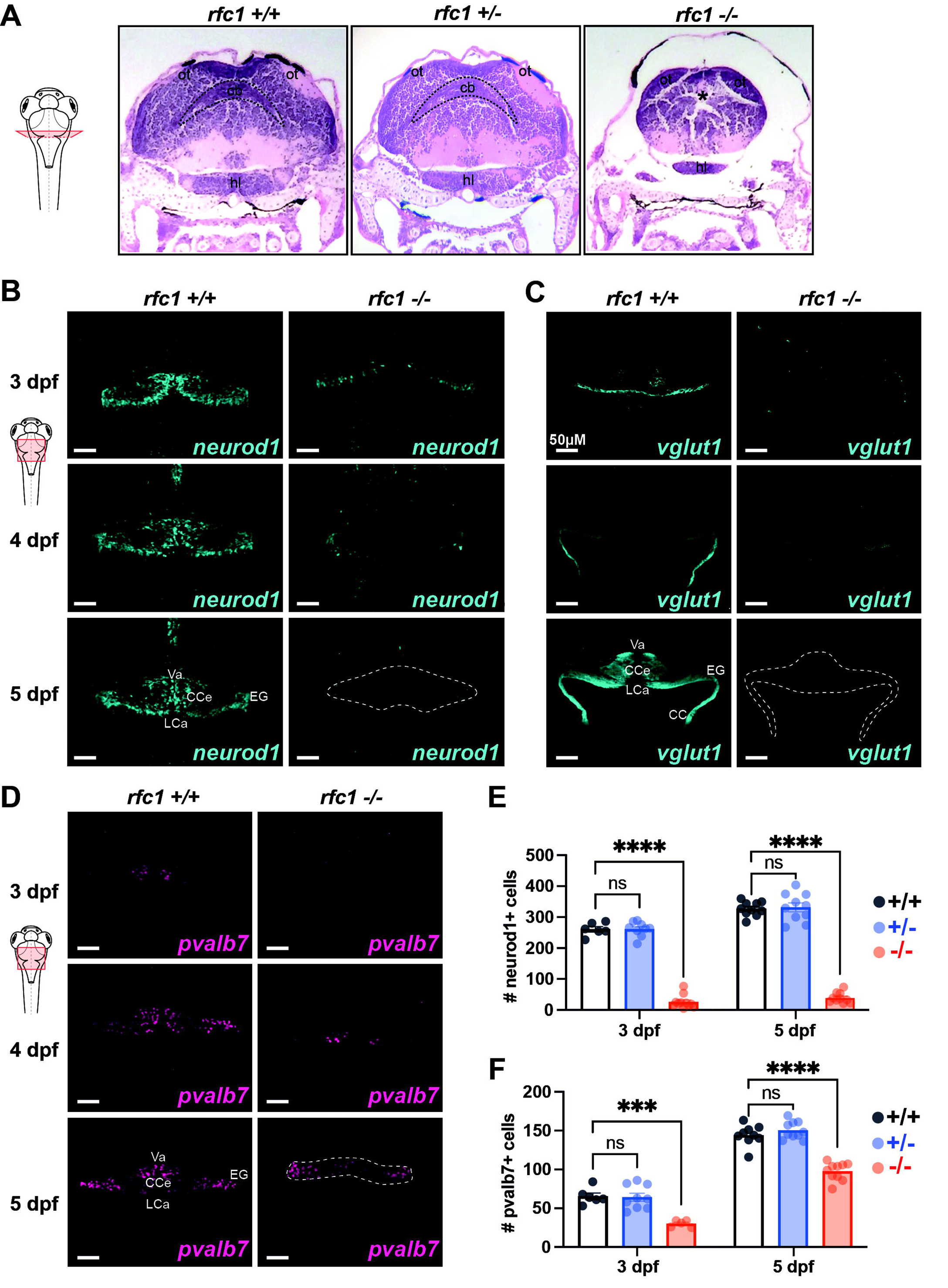
*Rfc1-/-* zebrafish larvae depict severe cerebellar defects. (A) Cross-section of 5 dpf zebrafish larva at the level of the *hindbrain and stained with hematoxylin and eosin. ot:optic tectum; cb: cerebellum; hl: hypothalamus.* (B) Immunolabelling using an anti-*neurod1* antibody to visualize granule cells in the developing cerebellum at 3, 4 and 5 dpf. The dotted line highlights the missing cerebellar structure at 5 dpf in *rfc1^-/-^* larvae. (C) Immunolabelling using an anti-*vglut1* antibody to visualize granule cells’ axonal projections in the developing cerebellum at 3, 4 and 5 dpf. The dotted line highlights the missing cerebellar structure at 5 dpf in *rfc1^-/-^* larvae. (D) Immunolabelling using an anti-*parvalbumin7 (pvalb7)* antibody to visualize Purkinje cells in the developing cerebellum at 3, 4 and 5 dpf. The dotted lines highlight the difference in structure at 5 dpf between *rfc1^+/+^* and *rfc1^-/-^* larvae. (E, F) Quantification of the number of neurod1-positive cells (E) or pvalb7-positive cells (F) at 3 and 5 dpf in +/+, +/- and -/- larvae. *ns : P > 0.05; * P* ≤ *0.05; ** P* ≤ *0.01; *** P* ≤ *0.001; **** P* ≤ *0.0001.* Va, valvula cerebelli; CC, crista cerebellaris; CCe, corpus cerebella; EG, eminentia granularis; LCa, lobus caudalis cerebelli.

### RFC1 is expressed in early neural progenitors and is necessary for their proliferation

To complement our previous observations and in order to dig further into RFC1-regulated brain cell populations, we performed deep RNA sequencing of single cells (scRNA-seq) from isolated larval brain of 4 dpf *rfc1-/-* and wildtype animals. We obtained the single-cell transcriptome of 16140 cells from wildtype and 6583 cells from *rfc1^-/-^* brains. We processed these data according to Seurat standard procedure of filtering and clustering to determine cell type identity based on genetic markers of zebrafish brain cell populations at relevant developmental timepoints ^38,39^. Consistent with our previous observations, we confirmed the general reduction in the number of cerebellar neuronal clusters defined as PCs (*pvalb7+, ca8+, aldoca+, ITPR1+*) and GCs (*neurod2+, neurod1+, zic5+, FAT2+, nebl+, olfm2b+, cbln12+, fabp4b+*) in *rfc1-/-* brains (Figure S4A-4B). Moreover, we found a significant loss of cell populations constituting clusters identified as neural progenitors (*her4.2+, her2+, sox2+*), cycling neural progenitors (*pcna+, mki67+, npm1a+*), neuronal progenitors (*dla+, dlb+, elavl3+, tubb5+*) and early neurons (*lhx9+, elavl3+, tubb5+*) (Figure 4A-4C and Figure S4D). Our findings suggest that early processes of neurodevelopment are perturbed in *rfc1-/-* fish, Given that cerebellar neurogenesis occurs as early as 2dpf ^40,41^, we next sought to perform scRNA-seq at 2 dpf. We dissociated the head of 2 dpf *rfc1^-/-^* and wildtype larvae and sequenced their individual transcriptomes (18632 cells from wildtype and 9805 cells from *rfc1^-/-^*). Although less severe than at 4 dpf, we noticed a significant reduction in the cellular density of the same clusters defined as neural progenitors, cycling neural progenitors, neuronal progenitors and early neurons (Figure 4D-4F and Figure S4C). Consistent with our prior expression analyses during neurodevelopment, we found that *rfc1* is mainly expressed in the neural progenitor and cycling neural progenitor clusters at 4 dpf and 2 dpf, respectively (Figure S4C-S4F). Altogether, these observations suggest that RFC1 regulates the development of early neural progenitors, particularly in the developing cerebellum. To confirm this hypothesis, we compared the expression level of specific markers of GC progenitors (GCp; *atoh1a, atoh1b, atoh1c*) ^42,43^, and of PCp (*ptf1a*) in *rfc1^-/-^* and wildtype larvae ^44,45^. We found a significant decrease in the expression of these markers in 3 dpf *rfc1^-/-^* larvae compared to their control siblings, both via RT-qPCR and whole mount *in situ* hybridization (Figure 4G-4K). We also showed that this reduction is sustained and progresses at later stages (Figure S4G-4J).

**Figure 4:**
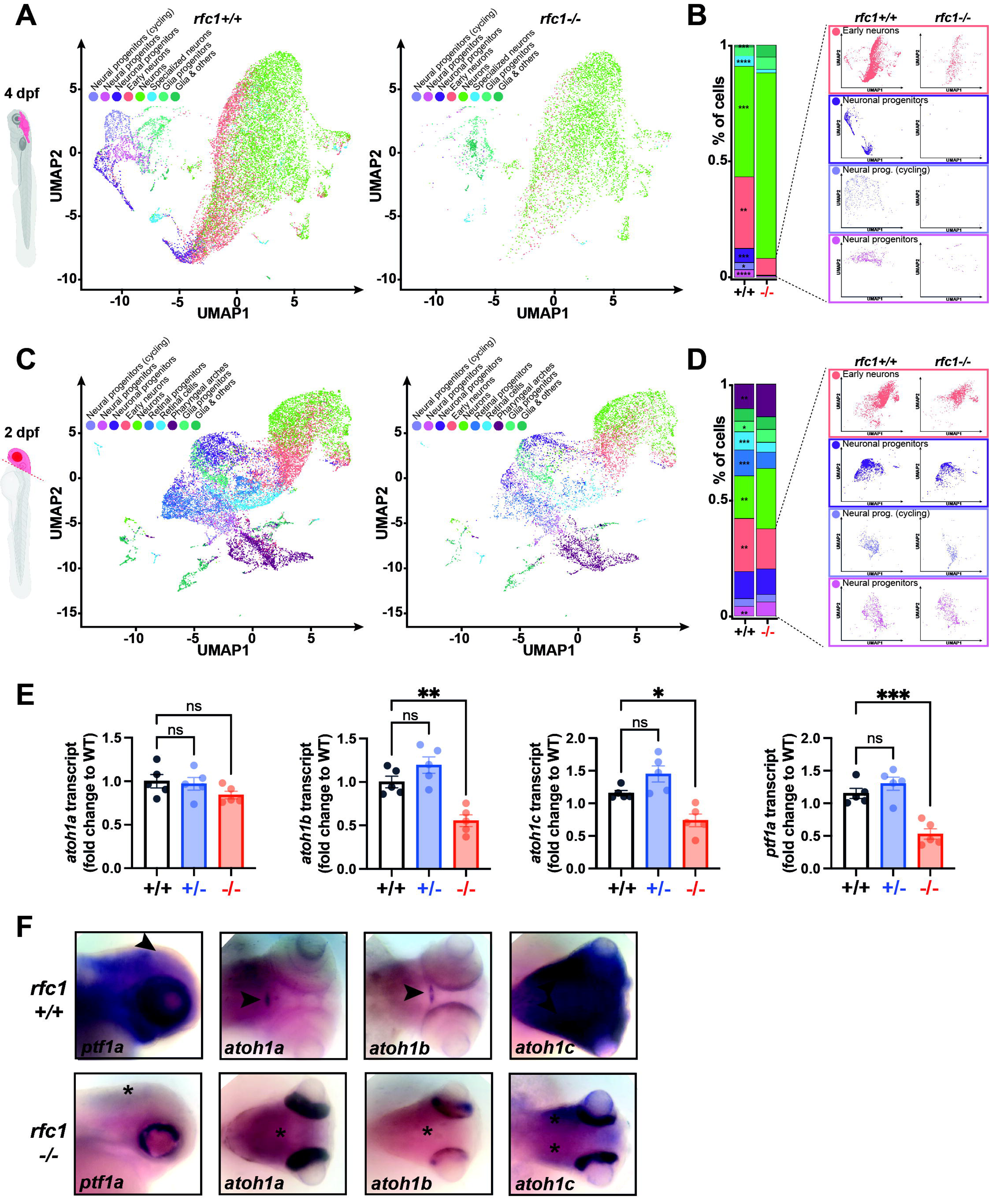
*RFC1* is expressed in early neural progenitors and is necessary for their proliferation. (A, C) Uniform manifold approximation and projections (UMAP) of single-cell RNA sequencing profiles from 4 dpf (A) and 2 dpf (C) *rfc1^+/+^* and *rfc1^-/-^*micro-dissected brains. Clusters of cells have been performed following Seurat standard procedure according to specific genetic markers (see Figure S4B, C). (B, D) Quantification of the percentage of cells among each cluster identified and individual UMAPs of clusters showing a significant reduction between *rfc1^+/+^* and *rfc1^-/-^*brains. (E) RT-qPCR of *atoh1a*, *atoh1b*, *atoh1c* and *pitf1a* on total RNAs extracted from whole 3 dpf +/+, +/- or -/- larvae. (F) Whole-mount in situ hybridization against *atoh1a*, *atoh1b*, *atoh1c* or *pitf1a* on 3dpf +/+ and -/- larvae. The expression in the wild-type cerebellum is shown with arrowheads and the missing expression in mutant larvae is shown with asterisks. *ns : P > 0.05; * P* ≤ *0.05; ** P* ≤ *0.01; *** P* ≤ *0.001; **** P* ≤ *0.0001*

Altogether, these results indicate that *RFC1* is expressed in early neural progenitors and is necessary for their development into differentiated neurons, particularly cerebellar neurons.

### RFC1 loss-of-function halts the proliferation of cerebellar neural progenitors and leads to their death

We next asked what molecular functions are regulated by RFC1 in neural progenitor cells in which it is expressed. To do so, we performed a differential expression analysis on our scRNAseq datasets at 2 and 4 dpf comparing *rfc1^-/-^*to wildtype sample (Figure 5A, 5B, Table S1 and S2). Within early progenitor clusters, we found multiple genes involved in early DNA damage response, cell cycle checkpoints, cell proliferation homeostasis, DNA repair and apoptosis to be differentially expressed between *rfc1^-/-^* and wildtype samples. In particular, we noted the overexpression of early DNA-damage sensor genes in neural progenitor clusters in *rfc1^-/-^* brains at 2 and 4 dpf (*mdc1*, *atm*, *atr*, *mre11* and Fanconi anemia complementation group genes (*fancd2*, *fancl*)). These genes are activated in response to DNA damage or replication blockage, and they halt cell cycle progression by controlling critical cell cycle regulators. Particularly, ATM and ATR are thought to be master controllers of cell cycle checkpoint signalling pathways that are required for cell response to DNA damage and genome stability. Consistently, we noticed an increased expression of many cell cycle checkpoints in the same mutant clusters such as *brca2*, *chek1/2*, *cdk6*, *cdk2* or *cdkn1a*. Notably, these genes are downstream DNA-damage triggers which stall cell cycle progression. Moreover, several genes involved in the negative regulation of cellular proliferation and cell cycle stalling, such as *nupr1a*, *yap1* and *btg2* ^46-48^, were found to be increased in mutant progenitor clusters. This is consistent with the decrease of proliferation markers such as *cenpf* or *cdca3* in these cell populations. We also found increased expression of multiple genes involved in DNA repair processes, such as *cenpx*, *xpc, ercc5/6* and *rpa1/2* Figure 5A, 5B). Those DNA repair and tumour-suppressor genes are activated by cell cycle checkpoint controllers upon alteration in DNA integrity ^49-53^. Finally, we showed that the expression of multiple pro-apoptotic genes (*tp53, baxa, casp8, mdm2, mdm4*) ^54-57^ and *p53* target genes (*rrm2, ccng1, rps37l sesn1, sesn3*) is increased in these same clusters (Figure 5A, 5B). Notably, these gene expression changes are predominant in neural progenitor clusters but also in retinal progenitors at 2 dpf and glia progenitors at 4 dpf.

**Figure 5:**
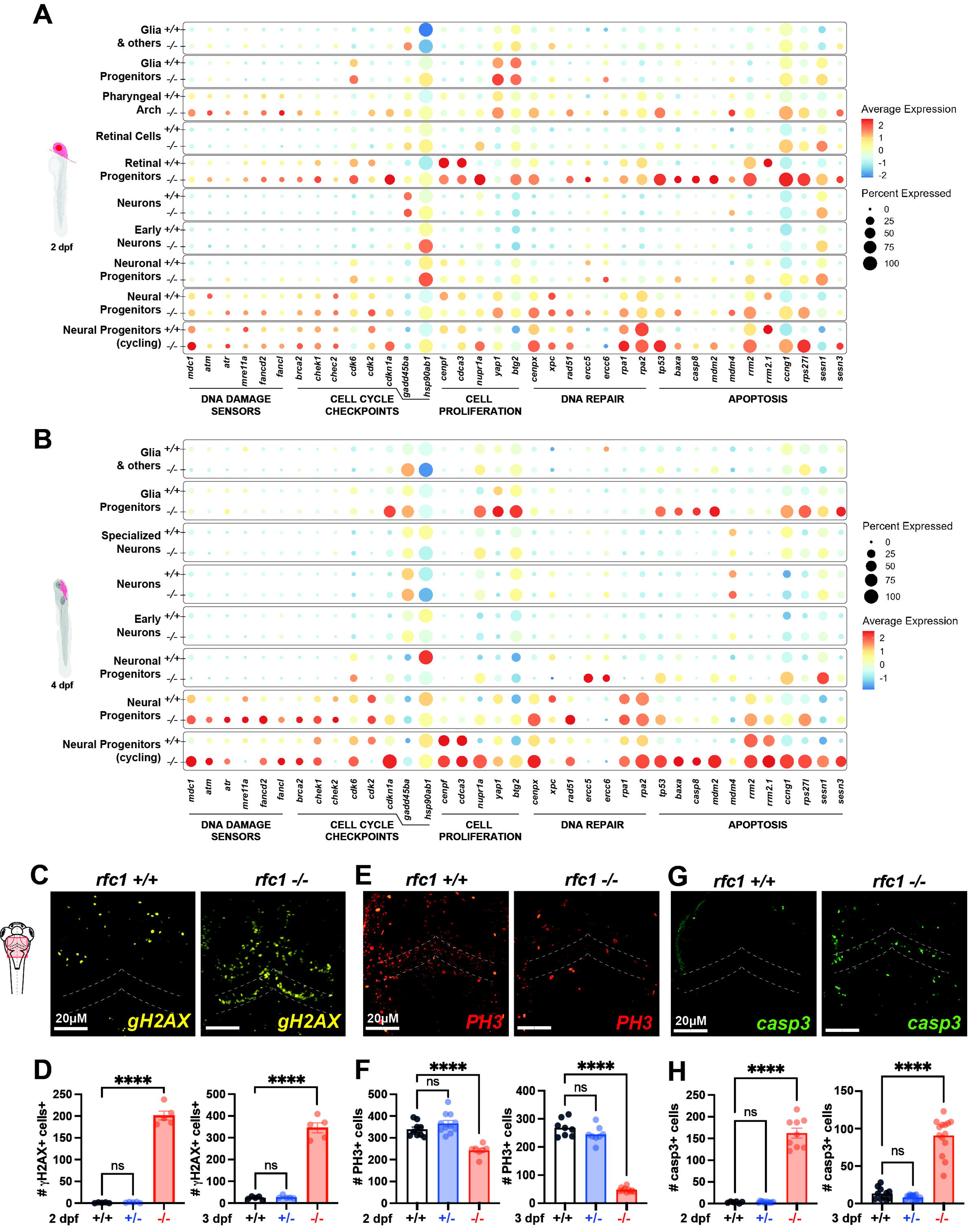
*RFC1* loss-of-function halts the proliferation of cerebellar neural progenitors and leads to their death. (A, B) Dotplots of differentially expressed gene expression between +/+ and -/- samples from individual scRNAseq clusters (see Figure S4B, C) at 2 dpf (A) and 4 dpf (B). (C) Immunolabelling using an anti-H2AX (a phosphorylated form of histone H2AX) antibody, to detect DNA damages (double-strand breaks) in the developing brain at 3 dpf. The dotted lines highlight the developing cerebellum. (D) Quantification of H2AX-positive cells in the developing brain of *rfc1^+/+^*and *rfc1^-/-^* larvae. (E) Immunolabelling using an anti-PH3 (a phosphorylated form of histone H3) antibody, a marker of mitosis in the developing brain at 3 dpf. The dotted lines highlight the developing cerebellum. (F) Quantification of PH3-positive cells in the developing brain of *rfc1^+/+^* and *rfc1^-/-^* larvae. (G) Immunolabelling using an anti-casp3 (activated-capsase3) antibody, a marker of apoptosis in the developing brain at 3 dpf. The dotted lines highlight the developing cerebellum. (H) Quantification of casp3-positive cells in the developing brain of *rfc1^+/+^* and *rfc1^-/-^*larvae. *ns : P > 0.05; * P* ≤ *0.05; ** P* ≤ *0.01; *** P* ≤ *0.001; **** P* ≤ *0.0001*.

We wanted to confirm these transcriptomic changes with a series of immunofluorescence labellings *in toto*. To do so, we first immunolabelled embryos using an antibody anti-phosphorylated histone (gH2AX), a marker of DNA damage at the level of the chromatin ^58^. We found a significant increase in the number of gH2AX-positive cells in the developing brain of *rfc1^-/-^*embryos compared to control siblings (Figure 5C, 5D). Moreover, we assessed cell proliferation using an anti phospho-histone 3 (PH3) antibody and showed that from 2 dpf, the number of proliferative cells in the developing hindbrain is significantly reduced in *rfc1^-/-^*embryos (Figure 5E, 5F). Notably, these changes are sustained at 3 dpf in *rfc1^-/-^* larvae compared to their siblings (Figures 5D, 5F). Finally, when quantifying the number of apoptotic cells at the same stages using an anti-activated caspase 3 antibody, we found a significant increase in the number of dying cells both at 2 and 3 dpf in *rfc1^-/-^* specifically (Figure 5G, 5H). Altogether, these observations suggest that *rfc1^-/-^* neural progenitor cells accumulate DNA damages, which trigger the stalling of cell cycle progression and, eventually, programmed cell death.

## Discussion

The *RFC1* gene has recently caught attention due to the recent identification of biallelic repeat expansions associated with a wide range of movement disorders, including late-onset cerebellar ataxias such as CANVAS ^14,16,17,19^. However, the underlying pathogenic mechanisms was unknown, and this lack of knowledge is largely explained by the fact that *RFC1* gene function has never been investigated *in vivo*. Our study is the first to investigate the function of *RFC1* during neurodevelopment, particularly in the cerebellum, the region primarily affected by ataxic cerebellar disorders. We showed that *RFC1* is expressed during cerebellar embryonic development, particularly in expanding neural progenitors in the early zebrafish embryo. Importantly, we showed that the loss of *RFC1* led to a severe cerebellar phenotype characterized by a significant reduction in PCs and GCs and a consistent motor dysfunction. We unveiled that the loss of *RFC1* jeopardizes the genomic integrity of these progenitor cells, which triggers cell cycle arrest and ultimately leads to their programmed death. Thus, our work is the first to show that *RFC1* plays a crucial role in cerebellar neurogenesis *in vivo*.

There are multiple DNA repair genes which have been associated with hereditary ataxias: *TDP1* is associated with spinocerebellar ataxia with axonal neuropathy-1 (SCAN1) ^59^; *XRCC1* is associated with autosomal recessive cerebellar ataxia ^60,61^; *PNKP* is associated with ataxia-oculomotor apraxia-4 (AOA4) ^62,63^; *APTX* is associated with ataxia-oculomotor apraxia-1 (AOA1); *PCNA* is associated with ataxia-telangiectasia–like disorder-2 (ATLD2) ^64^; *MRE11* is associated with ataxia-telangiectasia–like disorder-1 (ATLD1) ^65,66^ and *ATM* is associated with ataxia-telangiectasia ^67^. The mechanism by which ataxia-associated DNA repair genes lead to preferential cerebellar dysfunction is unknown. It has been suggested that the cerebellum and its main functional components, Purkinje cells, are particularly vulnerable to DNA damage, and are therefore more sensitive to changes in these proteins’ function. In this work, we showed that the expression pattern of RFC1 is not ubiquitous during embryo development, as it could have been expected. Moreover, in the postnatal cerebellum, RFC1 expression remains restricted to Purkinje Cells in the cerebellar cortex in mice (Figure 1B-E). Thus, this restricted and specific expression pattern could explain how deleterious genetic variations in such universal genes can lead to tissue-specific defects. As a result, instead of explaining the cerebellar specific effects of these variants by a greater sensitivity of this tissue /cells to DNA damage, it is possible that these genes are expressed in a very specific way at the tissue and cellular level, which explains the specificity of the defects when altered. Because they are involved in late-onset disorders, the mechanisms by which genetic variations in these DNA repair genes, such as RFC1, are mainly speculated in the context of post-mitotic neurons. However, our work suggests that these genes can be specifically expressed in the developing cerebellum and that they are essential for normal cerebellar neurogenesis. Notably, the investigation of these causative DNA repair gene’s function *in vivo* is hampered by the embryonic lethality of mouse KO models ^68^. Our study reveals that it could be insightful to investigate their function in vivo in other vertebrate species, such as zebrafish.

Our data showed that the developing cerebellum is primarily affected by *RFC1* loss-of-function, which is consistent with RFC1’s expression pattern in the developing hindbrain. However, it is worth noting that *RFC1* is also expressed in the developing eye, particularly at the level of the ciliary marginal zone (Figure 1A). This suggests that it plays a role in retinal neurogenesis and explains why the main transcriptomic changes we describe within brain progenitor populations are also found in retinal progenitor cells (Figure 5A, 5B). These results are also consistent with the striking reduced eye size phenotype observed in *rfc1^-/-^* larvae compared to their siblings. Although no retinal syndrome has previously been associated with *RFC1* hypomorphism, our data suggest that genetic variants in *RFC1* could be associated with it. However, understanding the mechanisms underlying this phenotype in the eye warrants future studies.

Our results can also be discussed regarding the uncertain pathogenic mechanisms underlying CANVAS. To date, no consensus exists on whether repeated expansions in *RFC1* can cause a gene loss of function nor a toxic gain of function. Indeed, although truncating variants in *RFC1* have been found in CANVAS patients ^20,21^ and that in silico analyses suggest repeat-formed secondary DNA structures which could prevent normal transcription ^69^, no irrefutable evidence of a loss of expression in *RFC1* has yet been provided in post-mortem tissues or in vitro cultures of patient cells. Nevertheless, our data provide evidence that RFC1 is essential for cerebellar development by ensuring genomic integrity during the expansion of neural progenitor populations. Thus, one can speculate that early deleterious changes in RFC1’s function or expression during development could affect the proper development of the cerebellum and that these developmental defects could also contribute, at least in part, to the pathogenicity of these repeat expansions. Thus, it would be very interesting to know if structural defects in cerebellar cortical organization could be present before the appearance of symptoms and contribute to the increased sensitivity of the system to DNA damage, for example. This would require generating relevant biological models of the disease, which are complex to mimic, particularly for large repeat expansions. Finally, our expression data in mice show that *RFC1* is expressed in post-mitotic calbindin-positive Purkinje cells at P60 in a mature cerebellum (Figure 5E). However, we noticed that RFC1 expression in mature PCs is mainly cytoplasmic. It is, therefore, possible to speculate that *RFC1* might play a distinct role than preserving the genomic integrity in these post-mitotic neurons. It is worth noting that other DNA replication and repair protein such as PCNA have been found cytoplasmic with a role in several cellular processes in differentiated cells ^70,71^.

In sum, our data improve our general knowledge of the *in vivo* function of the *RFC1* gene and open new avenues of study that may help elucidate the pathogenic mechanisms underlying so-called *RFC1*-related disorders.

## Limitations of study

Late-onset ataxia is, by definition, a disorder occurring at an adult stage. Therefore, our results show that the role of RFC1 during cerebellum neurogenesis cannot be directly transposed to explain the pathogenic mechanisms underlying CANVAS and other late-onset RFC1-related disorders. More work is needed to determine the function of RFC1 in post-mitotic cerebellar neurons, particularly for regulating the maintenance of genomic integrity against DNA damages against which the cerebellum seems particularly sensitive. Moreover, our functional investigation was carried out on zebrafish, which is a vertebrate. While the cellular organization of the cerebellum has been well conserved from teleosts to mammals, there are some obvious anatomical differences. For instance, the equivalents of the mammalian deep cerebellar nuclei reside in isolated eurydendroid cells located within the cerebellar cortex in teleosts ^41^. Nevertheless, the molecular pathway regulating its neurogenesis and maturation (including GC internal migration) are well conserved. Moreover, the cerebellum’s neuronal cell types, connectivity, and function are highly conserved from teleosts to humans, making the cerebellum one of the highest evolutionary conserved tissues throughout jawed vertebrate brains.

## Acknowledgments

We thank Pr Masahiko Hibi (Tokyo, Japan) for providing antibodies against pvalb7, neurod1 and vglut1 and associated protocols. We thank the CRCHUM animal facility, particularly the zebrafish platform, for their help and support in maintaining our fish colonies. We also thank Aurélie Cleret-Buhot from the CRCHUM imaging platform for her help in image acquisition. Finally, we thank the IRCM genomic platform for its rigorous work preparing single-cell RNA libraries. This work was funded by the Centre d’excellence en recherche sur les maladies orphelines – Fondation Courtois (CERMO-FC), by Génome Quebec and Ataxie Canada. E.S and S.A.P are supported by the Natural Science and Engineering Research Council (NSERC), Canadian Institutes of Health Research (CIHR), and are FRQS research scholars. M.T is a Junior 2 FRQS research scholar. F.N. received M.Sc scholarships from CRCHUM and the Faculty of Medicine/Université de Montreal. S.A received doctoral scholarships from CRCHUM, Faculty of Medicine/Université de Montreal, CERMO-FC, FRQS and CIHR.

## Author contributions

Conceptualization, E.S., S.A.P. and F.N.; Methodology: E.S., S.A.P. and F.N.; Investigation: F.N., S.A., S.T and C.Z.; Formal Analysis and Data Curation: S.A.; Resources: N.P, M.T, E.S; Writing Original Draft: E.S. and F.N.; Writing – Review & Editing: S.A.P., N.P. and M.T.; Visualization: N.P., E.S. and S.A; Supervision: E.S., S.A.P., N.P. and M.T.; Project Administration: E.S.; Funding acquisition: E.S.

## Supplemental information

Document S1. Figures S1–S4; Table S1 and S2.

Table S1. Excel file of all differentially expressed genes in individual clusters at 2 dpf, related to Figure 5A and 5B.

Table S2. Excel file of all differentially expressed genes in individual clusters at 4 dpf, related to Figure 5A and 5B.

## STAR Methods

### RESOURCE AVAILABILITY

#### Lead contact

Further information and requests for resources and reagents should be directed to and will be fulfilled by the lead contact, Dr. Éric Samarut

#### Materials availability

The rfc1-KO zebrafish line generated in this study is cryopreserved at the University of Montreal Hospital Research Center (CRCHUM) and is available upon request and signature of a Material Transfer Agreement between the CRCHUM and the requestor.

#### Data and code availability

- Single-cell RNA-seq data have been deposited at NCBI and are publicly available as of the date of publication. Accession numbers are listed in the key resources table. Original in situ hybridization images and microscopy data reported in this paper can be shared by the lead contact upon request.
- Any additional information required to reanalyze the data reported in this paper is available from the lead contact upon request.

### EXPERIMENTAL MODEL AND STUDY PARTICIPANT DETAILS

All experimental procedures involving mice were approved by the institutional ethics committee of the Université du Québec à Montréal (CIPA # 650) in accordance with the biomedical research guidelines of the Canadian Council of Animal Care (CCAC).. FVB/N mice initially purchased from Charles River (strain code 207) were maintained and bred in individually ventilated cages within the conventional animal facility of the Université du Québec à Montréal, under 12 h light–12 h dark cycles (7AM to 7PM) and with ad libitum access to regular chow diet (Charles River Rodent Diet #5075, Cargill Animal Nutrition). Embryos were generated by natural mating and staged by considering noon of the day of vaginal plug detection as e0.5. The brain was collected from embryos, pre-weaned juvenile (postnatal day P20) and weaned juveniles (P60) animals.

Routine zebrafish (*Danio rerio*) maintenance was performed following standard procedures (Westerfield) at 28.5 C under 12/12 hr light/dark cycles at the animal facility of the University of Montreal Hospital Research Center (CRCHUM), Montreal, QC, Canada. All experiments complied with the guidelines of the Canadian Council for Animal Care. The primary model used for this study was *rfc1* loss of function mutants. For generation of the *rfc1* loss of function model, zebrafish optimized Cas9 mRNA was synthesized using the mMESSAGE mMACHINE SP6 kit (Ambion) and a pCS2-nCas9n (addgne #47929) template linearized with NotI. A single-guideRNA sequence (5’-TGGGGTTGGTGCCACCTTAG-3’) targeting the 5th exon of *rfc1* was designed using the online tool CRISPRscan and ordered from Synthego, CA, USA. (Tubingen long fin (TL) wild-type embryos were collected for microinjection. A 1nL drop of a mix of 100 ng/µL of Cas9 mRNA and 30 ng/µL of gRNA was injected into one-cell stage embryos using a Picospritzer III pressure ejector. Genotyping of rfc1+/+ (wild-type), +/ (heterozygous) and -/- (homozygous) animals was performed by high resolution melting analysis (HRM) using genomic DNA extracted by boiling the embryo/larva/clipped caudal fin in 50 mM NaOH for 10 min and then neutralizing it with 100 mM Tris HCl (pH 8). HRM primers were designed using the Universal Probe Library Assay Design Center (Fwd: 5’ GCCATTACTGATGTGGGCATCTG 3’ and Rev: 5’ TGAAAGTCTCCTCCAGCAAATCC 3’). All primer sets are available upon request. The PCR reactions were made with 5 µL of the Precision Melt Supermix for HRM analysis (Bio-Rad #172-5112), 0.5 µL of each primer (10 µM) and 2 µL of genomic DNA and water up to 10 µL. The PCR was performed in a LightCycler 96 Instrument (Roche) using white 96-well plates. Two-step Evagreen PCR reaction protocol was 95*°*C for 2 min, then 45 cycles of 95*°*C for 10s and 60*°*C for 30s, followed by 95*°*C for 30s, 60*°*C for 60s, the temperature was increased by 0.02*°*C /sec until 95*°*C for 10 s, then cooling at 40*°*C. Curves were analyzed using the Roche LightCycler 96 software. Moreover, Polymerase Chain Reaction (PCR) and sequencing were used to verify the mutation’s exact sequence. The following primers were used for genomic PCR: Fwd 5′ TCACGCCTGTAATCCCAG CATTG 3′ and Rev 5′ TCTTGAAGAATAGCTGTGTTGTCCTGTCAC 3′. PCR amplicons were loaded on an agarose gel 1.5 % (A87-500G, FroggaBio) and single-band samples were sent to CES Génome Québec (Montréal, Canada) for Sanger sequencing.

To eliminate any putative off-target mutations, F0 founder was outcrossed with Tubingen long fin wild-type fish for at least three generations before phenotyping the embryos. All experiments were performed on larvae, and zebrafish sex is not yet determined at this stage. rfc1+/-fish were incrossed to obtain rfc1+/+, +/- and -/- from the same cross for experiments. Rfc1+/-fish of different generations have been used to verify the consistency of the phenotype.

### METHOD DETAILS

#### Morphological analysis

Rfc1+/+, +/- and -/- larvae were anesthetized buffered tricaine methanesulfonate (MS222) and dorsal and lateral images of the whole body were acquired using a 4x magnification under a stereomicroscope (Leica). Body length, eye size, head size and surface area were measured manually using ImageJ software (version 2.3.0). Body length was measured from the anterior tip of the head and the posterior top of the tail. Eye size and otic vesicle diameter or surface area were measured by specifying the eye and vesicle boundaries.

#### Survival monitoring

About 50 embryos were raised in a dedicated glass beaker, and death was monitored daily. Dead larvae were collected and processed for genomic DNA extraction and High-Resolution Melting genotyping on the same day. At least two independent batches of larvae were monitored, each containing at least 50 larvae. Data was gathered in GraphPad Prism (version 10) for Kaplan-Meier survival analysis.

#### Locomotion behavior

At 4 dpf, larvae were separated into single wells of a 96-well plate containing approximately 200µL of E3 medium. The plate was placed inside a Daniovision^®^ recording chamber (Noldus). Before the start of the experiment, larvae were habituated for 30 min in the dark. Larval locomotor activity was monitored over 1h light, 1h dark cycles over 4 hours. The Ethovision XT13 software (Noldus) was used to quantify the total swimming distance per period. Morevoer, the swimming tracks were extracted from the integrated view panel of the Ethovision software.

#### Mouse perfusion and brain cryosection

Mice brains were removed after either head decapitation (P0) or after isoflurane anesthesia and transcardic perfusion (P7, P11 and P60). Transcardiac perfusion was performed with 1X phosphate-buffered saline (PBS), followed by 4% (w/v) paraformaldehyde solution (PFA). PFA is commonly used to remove blood and preserve the brain for immunostaining. Removed mouse brains were postfixed in 4% PFA solution and left for 24 hours at 4°C. The PFA solution was replaced the next day with a 30% (w/v) sucrose solution (cryoprotection), and the brains were stored at 4°C until they sank to the bottom of the tube. A cryostat (LEICA CM 1950) was used to slice brain tissues into ten-micrometer sagittal brain sections (on glass slides) that were kept at -80°C.

#### Mouse brain Immunostaining

Brain slices on slides were permeabilized for 2h in a blocking solution (10% fetal bovine serum and 1% Triton X-100 in 1X PBS). Sections were then incubated overnight at 4°C with specific primary antibodies: anti-Calbindin D-28K (cb300, Swant, Switzerland; 1:200) or anti-PAX6 (Biolegand, UK; 1: 200). Subsequently, brain sections were incubated with secondary antibodies, diluted in the same blocking buffer, at room temperature for 2h. All sections were mounted and counterstained with DAPI (1:1000). A Nikon A1 confocal microscope equipped with NIS-Elements C software (Nikon) was used to acquire images, which were then analyzed using ImageJ.

#### Zebrafish In toto Immunofluorescent staining

Larvae were fixed in 4% paraformaldehyde (28908, Thermoscientific) overnight at 4 °C. The day after, PBS1X-Triton 1 % (X100-500ML, Sigma-Aldrich) was used for several washes over 1h. Larvae were then incubated in a blocking solution (2% NGS, 1% BSA, 1% DMSO, PBS1X-Triton 1 %) (005-000-121, Jackson Laboratory) for 30 min, and then rinsed several times with PBS1X-Triton 1%. Larvae were incubated in primary antibodies: anti-pvalb7 (gift from M. Hibi; 1:1000), anti-neurod1 (gift from M. Hibi; 1:500), anti-vglut1 (gift from M. Hibi; 1:1000) anti-gH2AX (GTX127342, GenTex; 1:1000), anti-PH3 (Millipore 09838; 1:500) or anti-cleaved caspase3 (Cell Signaling; 1:200). The following day, after several washes in PBS1X-Tween0.1% (P5927, Sigma), larvae were incubated with a secondary antibody diluted in blocking solution: goat anti-mouse Alexa Fluor 488 (Millipore Sigma A11029; 1:500), goat anti-rabbit Alexa Fluor 488 (Millipore Sigma A11034; 1:500). For imaging, larvae were positioned in 0.5% low-melting point agarose (16520-050, ThermoScientific).

and acquired using a Zeiss or an Olympus confocal microscope at a 20X or 40X magnification. Images were processed using ZEN (Zeiss) or Volocity (Quorum Technologies) software. Cell counting was performed manually using Image J.

#### Hematoxylin & Eosin coloration

Briefly, whole 4dpf larvae or microdissected adult brains were embedded in paraffin and sectioned transversally at 5 µm using Leica microtome (Jung Biocut). The hematoxylin staining was incubated for 4 min, washed with alcohol-acid and rinsed with tap water.

Finally, slides were incubated for 10s in lithium carbonate solution and for 2min in Eosin staining. Stained samples were mounted under a coverslip using Permount mounting media (3989, Sigma-Aldrich).

#### Probe cloning and in situ hybridization

A Specific 504bp rfc1 probe was cloned from 2dpf zebrafish cDNAs, using the following primers: Fwd-5’ TCACGCCTGTAATCCCAGCATTG 3’; Rev-5’ TCTTGAAGAATAGCTGTGTTGTCCTGTCAC 3’) within the pCS2+ vector using TOPO TA cloning kit (Invitrogen). Digoxygenin(DIG)-labelled sense and anti-sense RNA probes were *in vitro* synthesized using the DIG RNA Labeling Kit (Millipore Sigma, Roche-11175033910).

Whole-mount *in situ* hybridization was performed as described previously. Briefly, larvae were previously fixated in PFA 4 % overnight at 4°C. Embryos were digested with proteinase K (AM2546, Invitrogen): 1 and 2 dpf embryos were incubated at 10 µg/mL for 20 and 30 minutes, respectively; 3, 4, and 5 dpf larvae were incubated at 20 µg/mL for 30, 35, and 40 minutes, respectively. After several washes of PBS1X-Tween 0.1 %, embryos were fixed with PFA 4 % for 20 min. A prehybridizing step was performed at 69°C for 4 hours in MH Buffer (50 % formaldehyde (FOR001.1, Bioshop), 0.1 % Tween, 5X SSC (880-040-LL, Multicell), 11.5 mM citric acid pH6, 50 µg/mL heparin and 500 µg/mL tRNA). DIG probes (11093274910, Sigma-Aldrich) were incubated in MH Buffer at 69°C overnight. The day after, several washes with a prewarm medium were performed. Then, embryos were blocked for 1-4h at room temperature in Bloking Buffer (2% sheep serum, 2mg/ml BSA, PBT). Preabsorbed anti-DIG antibody was added overnight at 4°C incubation under gentle agitation. On the third day, samples were washed with PBS1X-Tween 0.1% and coloured using BCIP/NBT (11681451001, Sigma-Aldrich). Larvae were imaged in 80 % glycerol (G5516-1L, Sigma).

#### RNA extraction and qPCR

RNA was isolated from pools of 5 larvae using the RNeasy Plus Mini Kit (74134, Qiagen), according to the manufacturer’s protocol. Briefly, 500 ng of total RNA was used for cDNA synthesis using the SuperScript®Vilo™ Kit (11754050, Invitrogen).

The cDNA was diluted 1:10 and used for qPCR with SYBR Green I Master (Roche) on a LightCycler 96 instrument. Polr2d and actinB were used as reference genes for normalization. Primers were designed using A plasmid Editor (ApE) and span different exons: *rfc1*(Forward) 5′ CCA CAG AGA AGA AAC ACC CAG TAA C 3′ and *rfc1*(Reverse) 5′ GCC GTT CCA TTC TCT TTA GGT TTG 3′; *atoh1a*(Forward) 5′ GAG GCG AAG AAT GCA CGG ATT G 3′ and *atoh1a* (Reverse) 5′ TAG TAA GTC GGA CAG GGC GTT G 3′; *atoh1b*(Forward) 5′ TGG GAG TAA AGC ATC TGT CAG TGG 3′ and *atoh1b*(Reverse) 5′ TCT CCA ATG AAG GAA TGA CGC TTC TC 3′; *atoh1c*(Forward) 5′ GCT AAC GCC CGA GAG AGG AG 3′ and *atoh1c*(Reverse) 5′ TCT GTG CCA TCT GAA GCG TCT C 3′; *ptf1a*(Forward) 5′ CCA CAC AGT GAC GCC TTA AAC C 3′ and *ptf1a*(Reverse) 5′ GAG AGT GTC CTG CGA GAG GAG 3′; *polr2d*(Forward) 5′ AAC GCA AAG TGG GAG ATG TG 3′ and *polr2d*(Reverse) 5′ AGC GTC TCT GCG TTC TCA A 3′); *actinb*(Forward) 5′ CCA GCT TTT CAG CCT CAC TT 3′ and *actinb* (Reverse) 5′ CGG CAA TTT CAT CAT CCA T 3′. The 2-ΔΔCT algorithm, was used to *analyze* the relative changes in gene expression between samples.

#### Single-cell dissociation

Larval brains were microdissected from 4 dpf pre-genotyped larvae using fine forceps in calcium-free Ringers’ solution. For 2dpf embryos, heads were cut off using sharp microblades. A total of 4 brains or 3 heads were pooled per sample and transferred into a 1.5 ml tube containing 250 µl of dissociation solution containing 0.02% papain (LS003124, Worthington), 0.02% DNase I (LK003170, Worthington), and 12 mg/ml L-cysteine into 5 mL of DMEM/F12 (11320-033, Gibco). Tissues were digested at 37°C for 10 min and slowly pipetted up and down 10 times, then incubated for an additional10 min followed by pipetting up and down 10-13 times. To stop the enzymatic digestion, 1 ml of washing solution (0,65 % glucose 45 %, 0,5 % HEPES 1 M (15630-080, Gibco), and 5 % FBS qsp 10 ml DPBS 1X (14040-141, Gibco)) was added. Samples were then centrifugated at 4 °C at 800xg for 5 min. The supernatant was discarded and 50 µl of washing solution was added to resuspend the pellet. The cell suspension is filtered out with a cell strainer (136800040, SP Bel-Art, Wayne, USA) to remove aggregates or duplets. Single-cell suspension and cell number was assessed on a Kova slide and cell mortality was assessed using Trypan blue. Fresh single-cell samples were sent to the Montreal Clinical Research Institute (IRCM) genomic platform. Libraries were prepared using the Chromium Single Cell 3’ reagent kits. Reads obtained from the 10X Genomics platform were processed using the 10X Genomics Cell Ranger v7.1.0 pipeline and Cloud Analysis^72^, which includes STAR v2.7.2a alignment^73^ against the Danio Rerio GRCz11 genome with the corresponding Ensembl annotations (release 110) ^74^.

#### Single-cell data analysis

The bioinformatics analysis was performed in R (v4.3.1) using packages centred around Seurat v5.0.0^75^. Quality control steps included cell scoring with PercentageFeatureSet for mitochondrial genes (^mt-), ribosomal genes (^rp[sl]), and hemoglobin genes (^hba.*$|^hbb.*$). Cells were removed when they had poor sequencing depth (features ≤ 300; counts ≤ 500), abnormally high mitochondrial content (≥ 20%)^76^, or significant hemoglobin content (≥ 1%). Potential multiplets were removed by filtering out cells with outlier sequencing depth (features ≥ 6000; counts ≥ 50000). Features not present in at least three cells were also excluded from the analysis.

The Seurat RNA assay data was then log-normalized with NormalizeData default parameters, and scaled across all remaining genes. Additionally, an SCT assay was generated using SCTransform (vst.flavor = v2), and mitochondrial content was regressed during the gene expression transformation (vars.to.regress)^77^.

### QUANTIFICATION AND STATISTICAL ANALYSIS

The quantification of morphological and locomotion parameters was performed using Danioscope and Ethovision softwares respectively (Noldus) (described above). All data acquisition and quantification was performed by experimenter blind to the genotype of the larva. Details of biological and technical replicates, statistical test used and significance are mentioned in the corresponding figure legend. All experiments were repeated on, at least, two independent batches of embryos (represented by N), and the total number of animals per genotype used in each experiment was represented by *n*. Statistical analysis was determined using an unpaired Student’s t-test for comparing two conditions or a one-way analysis of variance (ANOVA) for comparing more than two conditions. The GraphPad PRISM software was used to create all the graphs, which are expressed as mean ± SEM. Significant thresholds were set at ns (p > 0.05); * (p <0.05); ** (p <0.01); *** (p < 0.001); **** (p < 0.0001). The GraphPad PRISM software was used to create all the graphs, which are expressed as mean ± standard error of the mean (SEM).

