## Supplementary Figure Legends for "*RFC1* regulates the expansion of neural progenitors in the developing zebrafish cerebellum"

**SUPPLEMENTARY INFORMATION**

Supplemental Table 1:

Supplemental Table 2:

**SUPPLEMENTAL FIGURE LEGENDS**

**Supplemental Figure 1: Protein alignment between human and zebrafish RFC1. Functional domains are highlighted in colours.**

**Supplemental Figure 2 : Morphological phenotype of rfc1-/- larvae compared to siblings.** (A) General morphology of *rfc1+/+, +/-* and *-/-* sibling larvae at 2, 3, 4 and 5 dpf. Scale bar 0.5 mm. (B) Quantification of morphological measurements.

**Supplemental Figure 3: *Rfc1-/-* zebrafish larvae depict severe cerebellar defects.** (A) Serial cross-sections of 5 dpf zebrafish larva at the hindbrain level and stained with hematoxylin and eosin. *ot: optic tectum; cb: cerebellum; hl: hypothalamus.* The lack of cerebellar structure in *rfc1^-/-^* larvae is indicated by asterisks. (B) Immunolabelling using an anti-*neurod1* antibody to visualize granule cells from a lateral view in the developing cerebellum at 3, 4 and 5 dpf. (C) Immunolabelling using an anti-*vglut1* antibody to visualize granule cells’ axonal projections from a lateral view in the developing cerebellum at 3, 4 and 5 dpf. (D) Immunolabelling using an anti-*parvalbumin7 (pvalb7)* antibody to visualize Purkinje cells from a lateral view in the developing cerebellum at 3, 4 and 5 dpf. *Va, valvula cerebelli; CC, crista cerebellaris; CCe, corpus cerebella; EG, eminentia granularis; LCa, lobus caudalis cerebelli.*

**Supplemental Figure 4:** ***RFC1* is expressed in early neural progenitors and is necessary for their proliferation.** (A) Individual uniform manifold approximation and projections (UMAP) of single-cell RNA sequencing profiles from 4 dpf *rfc1^+/+^* and *rfc1^-/-^* micro-dissected brains, showing clusters of cells marked as granule cells and Purkinje cells. (B, C) Clusters of cells have been performed following Seurat standard procedure according to specific genetic markers at 2 dpf (B) and 4 dpf (C). (D) UMAP of progenitor clusters at 2 and 4 dpf showing the expression of *rfc1* in these cells. (E) RT-qPCR of *atoh1a*, *atoh1b*, *atoh1c* and *pitf1a* on total RNAs extracted from whole 4 dpf +/+, +/- or -/- larvae. *ns : P > 0.05; * P ≤ 0.05; ** P ≤ 0.01; *** P ≤ 0.001; **** P ≤ 0.0001*
