## Supplementary figures and images for "*RFC1* regulates the expansion of neural progenitors in the developing zebrafish cerebellum"

### Figure S1

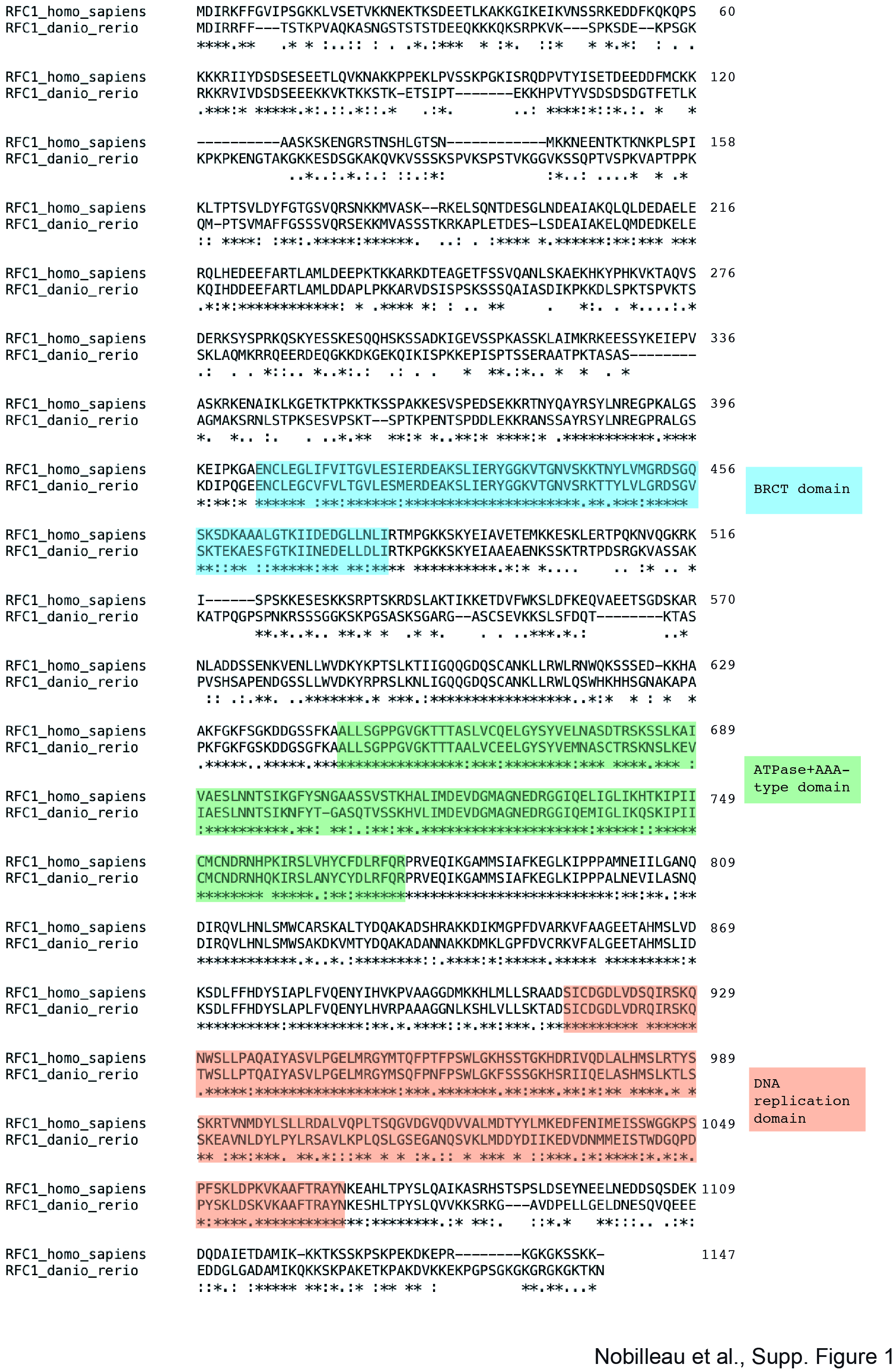

### Figure S2

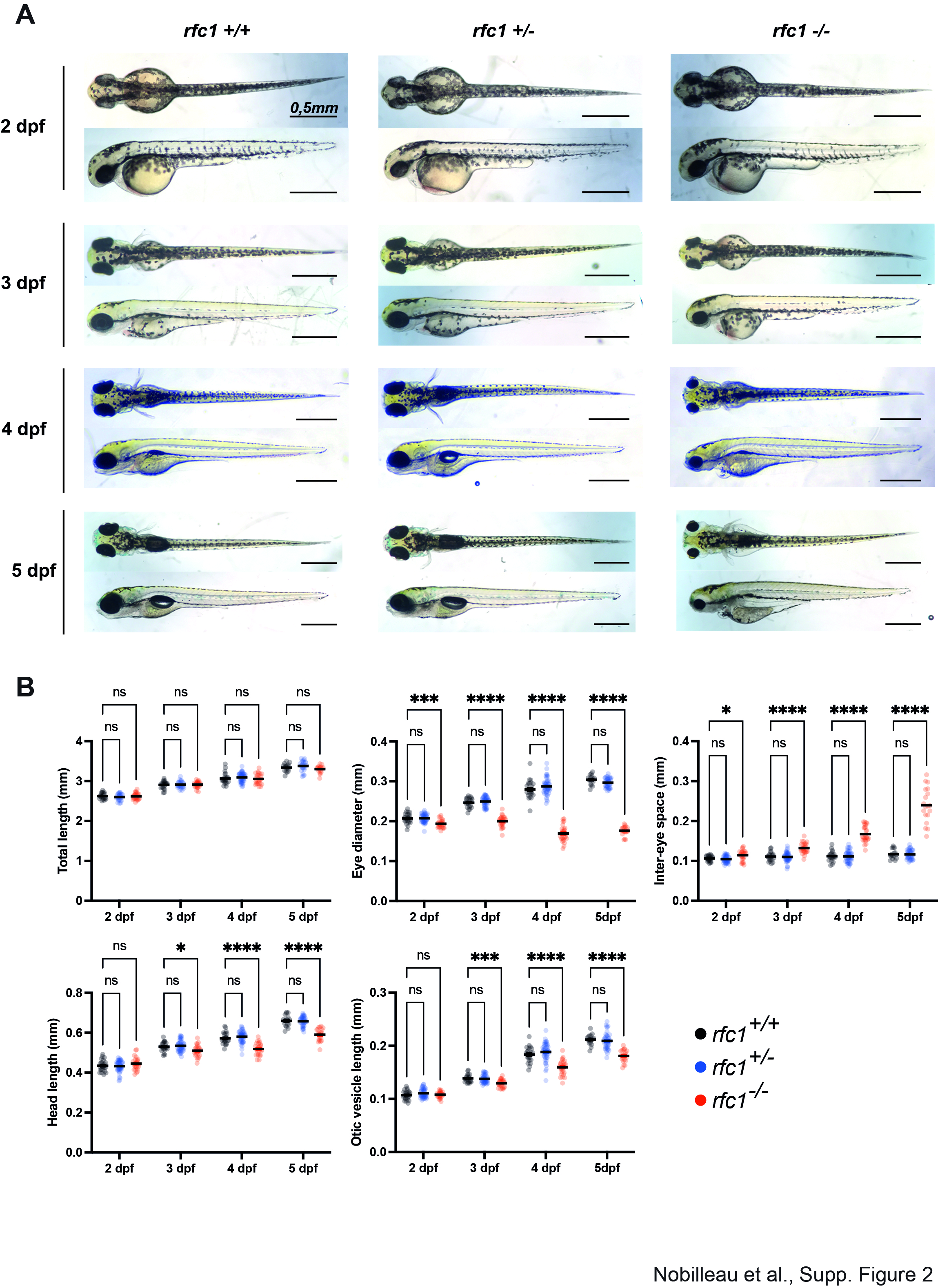

### Figure S3

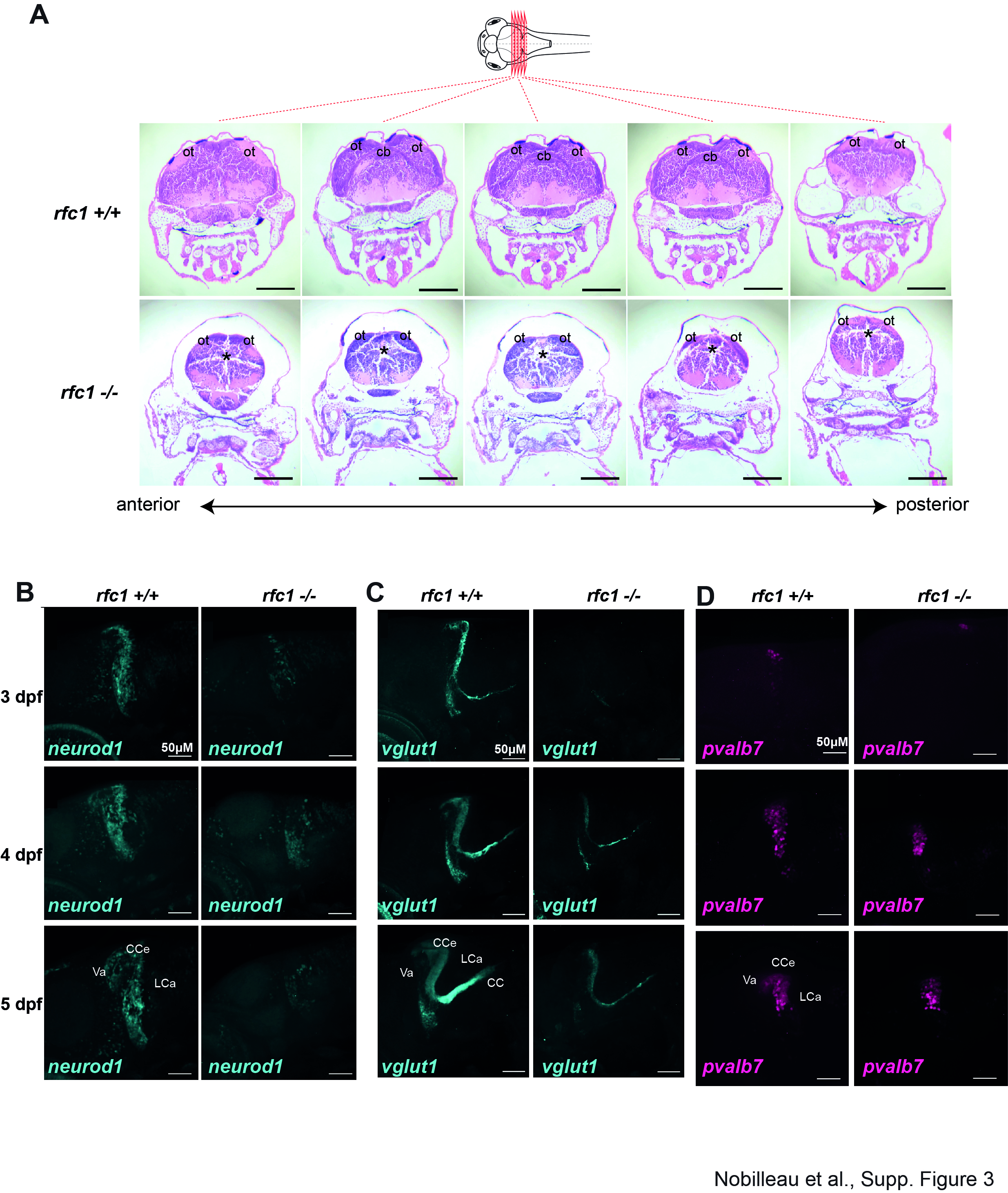

### Figure S4

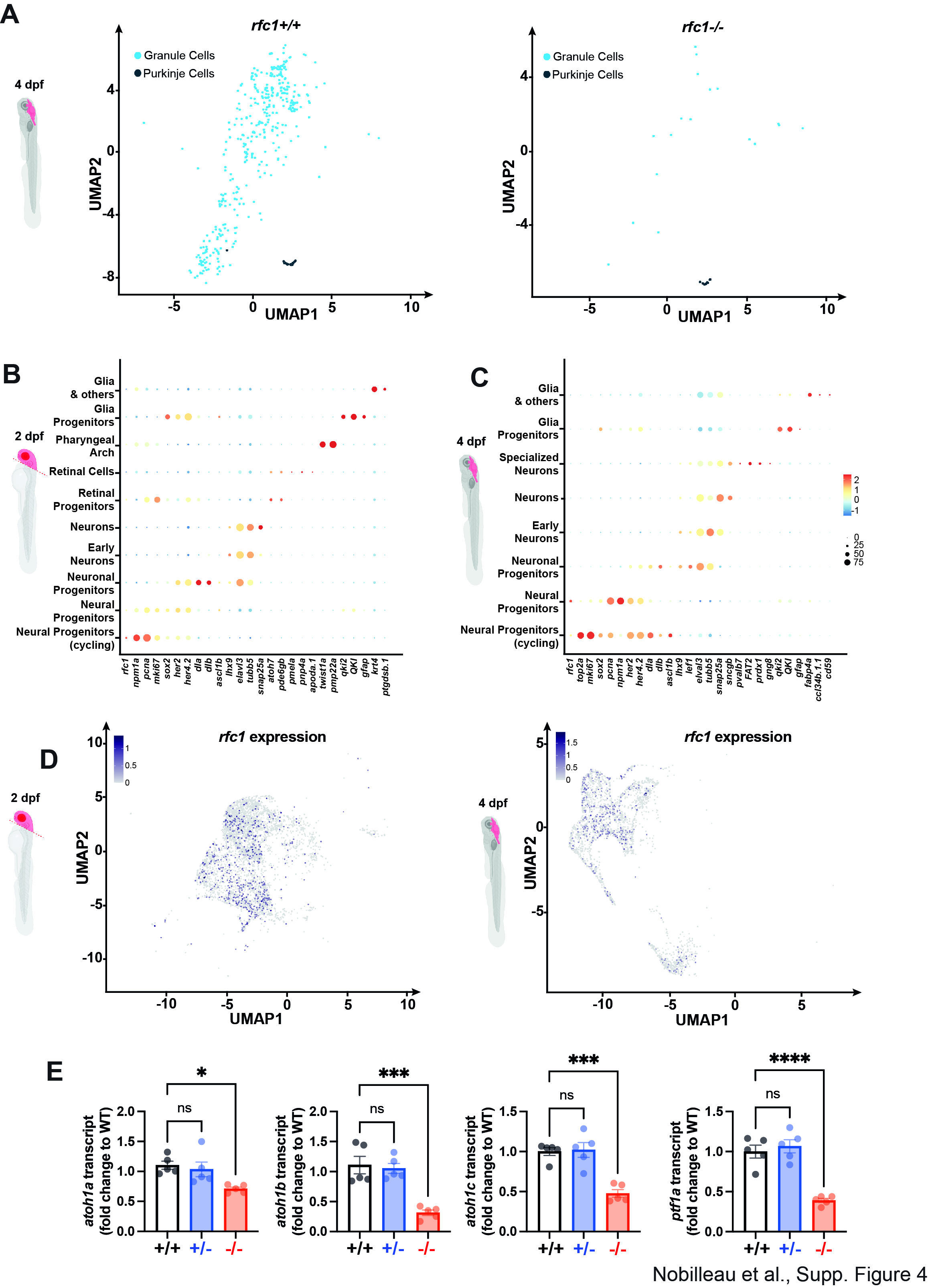
